# Asynchronous structural rearrangements govern constriction of the human nuclear pore complex

**DOI:** 10.64898/2026.07.31.742064

**Authors:** Erdong Ding, Matheus F. Mello, Vinícius G. Contessoto, Ronaldo J. Oliveira, Paul C. Whitford, José N. Onuchic

## Abstract

The nuclear pore complex (NPC) is responsible for transport between the cytoplasm and nucleus. CryoET experiments have resolved multiple conformations of the pore that are dilated or constricted. Since the dilation state will impact transport properties, there is much interest in understanding the energetic factors that regulate dilation/constriction. To this end, we applied a structure-based (Gō-like) model to study the mechanistic properties of large-scale constriction ( ∼ 100Å) in the human NPC. Even though the structures are symmetric, the simulations implicate asynchronous rearrangements during constriction. By considering variants of the model, we further characterize the relative contributions of individual proteins to this collective process. Together, this provides an energetic/structural foundation that can guide development of precise physical approaches for studying this elusive motion.

## I. INTRODUCTION

Eukaryotic organisms differ from bacteria in that each cell contains a nucleus encapsulated by a nuclear envelope, which separates the cytoplasm and nucleus. Consequently, large molecules that are synthesized in the nucleus, such as mRNAs, cannot diffuse passively through the nuclear envelope. Instead, transport is enabled by the nuclear pore complex (NPC).^1–4^

The human NPC is a massive macromolecular assembly embedded in the nuclear envelope (Fig. 1a).^1–8^ In humans, the NPC is composed of eight subunits (spokes), each containing over 100 proteins (nucleoporins, or NUPs). These sub-units adopt a spoke-like arrangement, together forming a ring-shaped structure with eight-fold rotational symmetry. Along its central axis, the NPC may be described in terms of three layers of rings, connected by Bridge NUPs (Fig. 1a). The rings facing the cytoplasm and the nucleus constitute the outer layers, and they are referred to as the cytoplasmic ring (CR) and the nuclear ring (NR). The inner layer is connected to the fused part of the nuclear membrane. The domain facing the pore is called the inner ring (IR), while the so-called luminal ring (LR) domain resides in the lumen formed by the inner and outer membranes of the nuclear envelope. Disordered protein tails extend from the IR to form a selective barrier within the pore that modulates nucleocytoplasmic transport.^7,8^

**FIG. 1.**
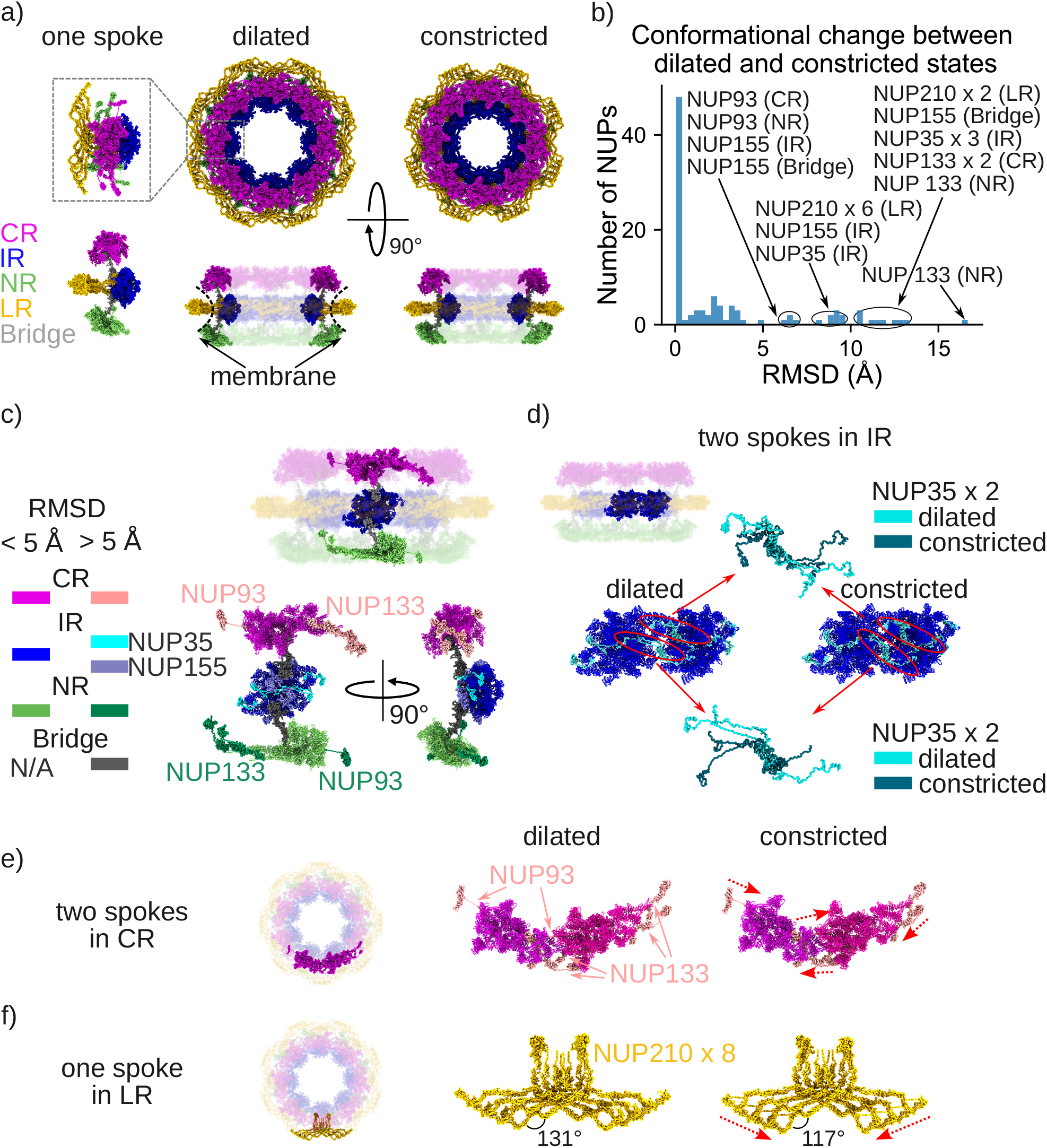
Structural comparison of the dilated and constricted states suggests possible candidates that contribute to NPC constriction. a) The dilated and constricted structures from top and side views. The side view shows two opposite spokes opaque and the other six transparent. The conformation of a single spoke is shown in the top and side views. Comparing the two states, the IR and LR undergo more obvious constriction than the CR and NR. b) RMSD distribution of all the NUPs between the dilated and constricted states. Most NUPs exhibit little conformational change between the two states, leaving those with an RMSD greater than 5 Å of particular interest. c) The structure of a single spoke of all the NUPs except the LR. The NUPs with an RMSD larger than 5 Å are highlighted in both the top and side views. Most highlighted NUPs are located at the interfaces between spokes or at the interfaces between different rings. d) The inter-spoke IR interface in the dilated and constricted states. The four linker NUPs (NUP35) form two dimeric structures at the inter-spoke IR interface, bridging the two spokes. Both dimeric structures exhibit a motion in which the internal angles decrease, facilitating the constriction. e) The inter-spoke CR interface in the dilated and constricted states. Three NUPs at the interfaces undergo constriction, where the ordered regions are pulled closer by altering the structure of the disordered region in between. The red arrows associated with the constricted structure indicate the constriction of NUP93 and NUP133 (colored pink). f) One spoke of the LR in the dilated and constricted states. The red arrows associated with the constricted structure indicate the LR bending motion accommodating the constriction.

It has only recently become possible to visualize the overall architecture of the NPC in living cells. Combining experimental tools from structural biology with physical modeling, structural descriptions of individual NUPs in the NPC have been constructed, which can be arranged into electron density maps of the complete assembly.^9–11^ Recent advances in imaging have produced electron density maps of the human NPC in isolated nuclear envelopes with a resolution of 12 Å, and *in cellulo* at 50 Å resolution. With the aid of previously determined structures of components, physical modeling and AI-based tools, over 90% of the human NPC structure has been reconstructed.^7^

In the past two decades, physical modeling on NPC systems has focused primarily on the properties of the transport barrier formed by the disordered proteins in the central channel of the nuclear pore^12–18^ or on individual NUPs^19^. However, global motions of the NPC were also explored with elastic network models.^20^ Benefiting from recent advances in imaging technologies and high-performance computing, physical models have been applied to investigate the NPC from a variety of perspectives, including the structural modeling^7,21,22^, transport kinetics and barrier dynamics within the central channel^23–28^, NPC disruption by the HIV capsid^29,30^ and NPC assembly dynamics^31,32^.

The human NPC exhibits a constricted conformation when isolated, and is dilated *in cellulo*, where external forces act on the nuclear membrane.^7,21^ While there has been notable progress in describing the structures of NPCs, the precise factors that govern constriction and dilation are not well understood. Change in diameter has been inferred to correlate with the rate of nucleocytoplasmic transport.^21,22,33^ Although the NPC itself does not have motor-like activity, evidence indicates that NPCs maintain larger diameters *in cellulo* due to their interactions with the nuclear envelope, which tends to expand in living cells, due to active mechanical processes.^21,22^ For example, under conditions of osmotic shock or energy depletion in *S. pombe* cells, the yeast NPCs exhibit smaller diameters that correlate with decreases in nuclear volume.^21^ This implies that interactions between the membrane and the NPC may be thought of in terms of an external potential that can ensure the NPC remains in an energetically strained dilated state in the cell. However, when the properties of these active cellular forces change, mechanical coupling between the NPC and the nuclear membrane leads to conformational rearrangements, including partial/total constriction.

In this work, we investigate the non-equilibrium dynamics associated with constriction in the human NPC. Specifically, we perform molecular dynamics simulations using a structure-based (SMOG) model^34–36^, which allows for the full constriction process to be simulated without introducing artificial steering/biasing terms. Our results reveal that, while minor local kinetic barriers exist within the NPC, the overall constriction is not rate limited by a single dominant free-energy barrier. Rather, the NPC appears to behave as an over-damped nonlinear spring, where the motions of individual domains can occur on disparate timescales. That is, the CR and NR contract an order of magnitude faster than the IR and LR. Furthermore, we show that this global transition is driven by large, coordinated conformational changes in only a few specific nucleoporins, such as the bending of NUP210 in the LR and structural rearrangement of NUP133 and NUP93 at the outer ring interfaces. Together, these simulations implicate the presence of geometric frustration within the NUP35 linkers at the inner ring hub, which may allow them to dynamically respond to external membrane forces.

## II. METHODS

### A. Structure refinement

The conformation of the constricted (PDB ID: 7R5K^7^) and dilated (PDB ID: 7R5J^7^) states from the PDB Bank could not be directly applied in the simulations because they both have misplaced residues. To model these residues, we first generated the all-atom structure based model (SBM) force field of two adjacent spokes for both constricted and dilated systems with the SMOG2 package^35,36^; the Amber variant of the all-atom SBM^37^ was used to generate the all-atom SBM force fields in this study. We then performed energy minimization for both systems and regenerated the force field with the relaxed structures. After minimization, a number of bond lengths differed by more than 10% of the Amber force field values^38^. All bonds with atypical lengths were associated with residue misplacement (*e*.*g*. forming knots or a side chain crossing through a ring). For these cases, we regenerated the local conformations of the residues using the Modeller package^39^, and the results were examined to ensure all the chemical bonds satisfy the 10% criterion discussed above.

The new conformations of two adjacent spokes were then processed through the same procedure, *i*.*e*. force field generation and energy minimization, to generate the C_*α*_ SBM. This step is to obtain the conformation of an asymmetric unit to build a whole human NPC complex by rotation. This asymmetrical unit was selected to have all the inter-spoke interfaces so that the interfaces between two subunits can be described accurately. Then, we froze the atoms not belonging to the asymmetric unit and not involved in inter-spoke interactions during the energy minimization process. The constricted and dilated states of the entire human NPC were then reconstructed using the corresponding constricted and dilated asymmetric units.

### B. SBM and molecular dynamics simulation

Based on energy landscape theory^40,41^, the SBM is a phenomenologically-guided Hamiltonian, where the native structure of a given system is its energetic minimum and the non-bonded energy is roughly a function of the reaction co-ordinate *Q*, a representation of structural nativeness. The C_*α*_ version of the SBM takes all the C_*α*_ atoms in the protein, and is constructed as follows,

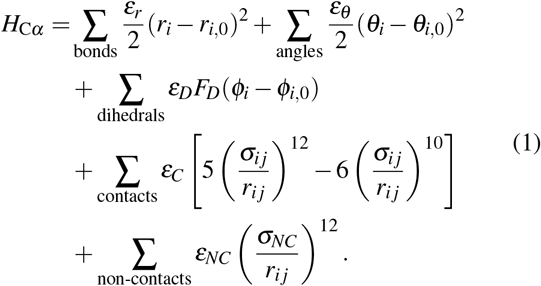

Here, the *r*_*i*,0_, *θ*_*i*,0_, *φ*_*i*,0_, and *σ*_*i j*_ are extracted from the native structure, all the energies are measured with reduced units (see below), and we have

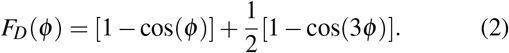

For the human NPC system (624040 C_*α*_ atoms), the Hamiltonians for both dilated and constricted states were generated with SMOG2 using the conformations obtained in the steps above. All parameters were set to the default values. Since the force field can lack exact symmetry due to limited precision allowed by the PDB format, the definitions of contacts were symmetrized after force field generation by manually appending the contacts missed in one spoke but present in any others. The systems were first equilibrated with the dilated-state force field at *T* = 0.6 (in reduced units, approximately corresponding to 300 K.^42^ See the next paragraph for details). Then, ten frames from the equilibrated ensemble were randomly selected and used as initial conformations for subsequent simulations. For each initial conformation, two independent simulations were performed with the constricted-state force field, giving a total of 20 simulated events. The simulations were performed with OpenSMOG v1.2^36^ and OpenMM v8.2^43^ using Langevin dynamics. Each simulation was performed for 8 × 10^7^ steps (4 × 10^4^ *t*_0_). For the perturbed Hamiltonians, 10 copies were simulated.

The reduced units used in SBMs warrants further explanation. Here, the energy unit used in the Lennard-Jones potential is denoted as *ε*_0_, and the energy minimum of all the particle-particle Lennard-Jones potentials in the SBM is set to be *− ε*_0_. The time unit, denoted as *t*_0_, is then derived from *ε*_0_. Although the exact value of *ε*_0_ can not be evaluated *a priori* for the phenomenological model, in practice, the energetic interactions in biological systems are known to be on the order of *k*_*B*_*T*, where *k*_*B*_ is the Boltzmann constant and *T* is the temperature.^44^ This means *ε*_0_ can be approximated as 1 kcal/mol, and the room temperature is roughly 0.6 *ε*_0_ in reduced unit, with *k*_*B*_ set to be 1. Since the coarse-grained model is associated with an accelerated diffusion coefficient, converting reduced units to an experimental unit requires additional calculations^45^. However, these types of comparisons have not been performed for the NPC, so we will report the results in terms of reduced time units.

### C. The reaction coordinate *Q*

*Q*, defined as the ratio between the number of formed native contacts and the total number of contacts that are defined in the model, can serve as a reaction coordinate for protein folding or conformational changes. The human NPC has more than 2 million native contacts. The majority of them are common between the constricted and dilated states, and they do not drive the kinetic process. Therefore, we have identified so-called “unique contacts” where the ratio of native distances between the constricted and dilated states exceeds a threshold, as described previously.^46^ *Q* is calculated for only the unique contacts throughout the manuscript.

For a kinetic process, we denoise the reaction coordinate *Q*. A typical contact undergoes a rapid formation process at time *t*_*C*_, which means *Q* of a typical contact is 0 before *t*_*C*_, goes to 1 at *t*_*C*_, and then occasionally drops to 0 while its average, or formation probability, ⟨*Q⟩* = *p* remains close to 1 after *t*_*C*_. If the formation probability *p* is less than 0.7 after *t*_*C*_, we consider the contacts to have a formation time larger than the simulation time. For the formed contacts, the fluctuations in *Q* introduce noise due to the large number of unique contacts. Thus, we constructed a monotonically increasing function to capture only the formation of the contacts as a denoised *Q*. For a given contact at any time *t*^*’*^, we calculate the average *Q* before and after *t*^*’*^, 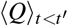 and 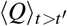. We can then estimate the formation time of the contact. For a typical contact, the value 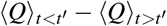 reaches its maximum at time *t*_*C*_, with 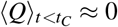 and 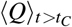 being the actual contact formation probability *p*. The denoised *Q*(*t*) is then defined as the fraction of contacts with *t*_*C*_ *< t*.

## III. RESULTS

### A. Large-scale NPC constriction is facilitated by a small number of NUPs

Inspection of available cryoET structures of the NPC suggests that a small number of proteins are likely to contribute to the dynamics of constriction. The constricted structure of human NPC (PDB ID 7R5K^7^) was obtained previously by fitting atomic models to a cryoET density of 12 Å obtained for the purified nuclear envelope *in vitro*. The structure of the dilated conformation (PDB ID 7R5J^7^) was constructed by first aligning the constricted structure to an ET density of 50 Å resolution, followed by flexible-fitting structural refinement protocols.^7^ While the levels of resolution introduce ambiguity into some of the structural details, the available models revealed the overall structural organization of both states, as well as implicated a range of local conformational changes that are associated with the ∼ 100 Å conformational transition.

Although human NPC is a massive complex that undergoes a large conformational rearrangement, most of its constituent proteins (referred to as NUPs) adopt similar conformations in the dilated and constricted states. However, there is a small group of NUPs that undergo substantial reorganization, which is the focus of the current study. To identify NUPs that are likely to influence constriction, we calculated the root mean square deviation (RMSD) between states for each NUP (Fig. 1b), and highlighted those with an RMSD greater than 5 Å (Fig. 1c). Based on this, we partition the various NUPs by their likelihood of contributing to constriction dynamics.

First, within the “linker-scaffold” structure in the IR spokes^8^, the conformations of most scaffold NUPs depend minimally on the constriction/dilation state (Fig. 1c), except for two copies of NUP155, possibly due to their interactions with the Bridges (Fig. 1c). On the other hand, the linker NUPs (NUP35), for which there are four per spoke, undergo large-scale changes (Fig. 1d). The linker NUPs form symmetric dimeric arrangements between adjacent spokes, which appear to have very flexible regions that may respond to the diameter change by altering the internal angles of the linkers (Fig. 1d). With regards to CR and NR, the major structural elements are similar, and we will use CR as a reference for the following discussion. The individual structures of CR NUPs largely remain similar between the constricted and dilated states. However, the NUP133 proteins are an exception, as they bridge the interface between adjacent spokes (Fig. 1c, e). Another NUP with a large RMSD is NUP93, also located at the inter-spoke interface. Together, these NUPs have structured regions that are connected by unstructured regions (Fig. 1c, e). They exhibit conformational changes in the unstructured regions that shorten the distances between structured units to accommodate the constricted structure (Fig. 1e). NUPs in LR (NUP210, Fig. 1f) and the Bridge NUPs (NUP155, Fig. 1c) between CR/NR and IR are also associated with large RMSD values between the two states. NUP210 undergoes a bending-like rearrangement that is associated with a decrease in pore diameter. Suggested by the analysis, NUP35 in IR, NUP93 and NUP133 in CR and NR, NUP155 in IR and Bridge, NUP210 in LR, are more likely to have an impact on the global constriction process.

### B. The human NPC constriction shows asynchronous timescales at the global level

While structural analysis can suggest which NUPs may contribute to dynamics, determining the ordering of conformational substeps and potential intermediates requires additional strategies. To this end, we used structure-based models (SBMs, or SMOG models)^35,36^. This class of models has been extensively applied to investigate the mechanistic and kinetic properties of large biomolecular systems, including the folding dynamics of complex protein structures^47^ and multi-domain proteins^48^, as well as partial unfolding/refolding processes associated with functional dynamics.^49,50^ With these models, we use molecular dynamics simulations to characterize the likely sequence of conformational motions during the transition from dilated to constricted states. Here, we use a C_*α*_ representation (624040 residues in total)^34^, where the constricted conformation (Fig. 2a) is defined to be the global minimum on the potential energy landscape. The residues that are in contact in the constricted conformation are explicitly defined to be attractive, while all others only interact through excluded volume. It is worth noting that these models may be extended to investigate the contributions of any number of factors, such as electrostatics^51,52^. However, by applying an electrostatics-free variant of the model, the principal objective is to determine whether there is a likely sequence of conformational substeps during constriction, and if structural features lead to sterically-induced free-energy barriers.

**FIG. 2.**
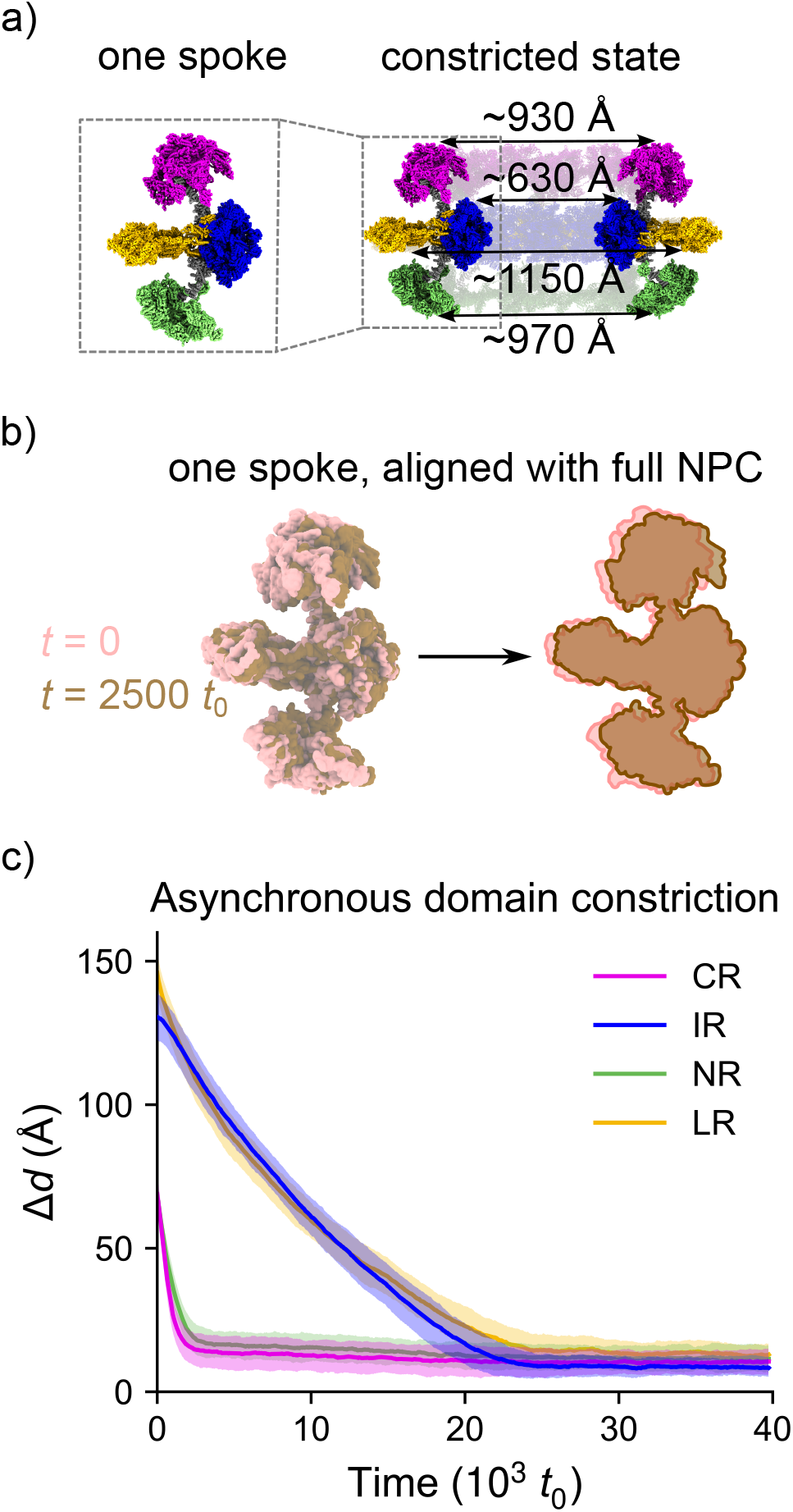
Constriction simulations show distinct timescales for different rings in human NPC. a) The constricted state of the human NPC in a cutaway view with the diameters of all the rings. The CR, IR, NR, LR and Bridge are colored magenta, blue, green, yellow and grey, respectively. The conformation of a single spoke is shown on the left for comparison with panel b). b) The conformations of a single spoke during a representative simulation at *t* = 0 and *t* = 2500 *t*_0_, obtained by first aligning the full NPC to the constricted state and then superposing the spoke. The early stage of NPC constriction features a rapid constriction of the CR and NR. c) Diameter changes of CR, IR, NR, and LR during the simulation. The CR and NR show a similar timescale of constriction, while the IR and LR exhibit a significantly larger timescale. The average IR diameter among spokes and simulation copies is plotted, where the shaded areas represent the standard deviation.

We performed 20 simulations from the dilated state to the constricted state. In each simulation, the system was first thermally equilibrated and then simulated until the constricted conformation was adopted. In the current study, we use multiple reaction coordinates to describe the overall characteristics of these transitions. First, the distances between the centers of mass of CR, IR, NR, and LR provide measures of the diameter *d*. Specifically, the distance between CR, IR, NR, or LR of opposing spokes (spoke *i* with *i*+4) yielded four distinct measures of diameter for each spoke pair. In addition, since the energetics in this model is predominantly associated with native contacts, it is natural to characterize the dynamics using the fraction of native contacts formed, denoted *Q*, as a reaction coordinate.^34,53–56^ When describing the dynamics along each coordinate, we report all time units in terms of the reduced unit *t*_0_, which may be converted to an experimental timescale.^45^ However, since the current study does not explicitly include membrane effects, the absolute timescales would not be intended to represent firm predictions. Instead, the main utility of comparing timescales is that they can be used to determine whether different conformational motions are likely to be strongly coupled, and whether a distinct ordering of events may arise from structural considerations.

The first observation in these simulations is that constriction does not appear to be associated with a large-scale free-energy barrier. That is, the folding or conformational change kinetics of large biomolecules can often be approximated to satisfy a Langevin equation and the corresponding Fokker-Planck equation in a lower-dimensional space, given that the reaction coordinate is appropriate.^57–62^ When using the NPC diameters to describe constriction, the effect from the noise term, or the diffusion term from the Fokker-Planck equation, is small (Fig. 2c). The diameters of different spoke pairs show similar dynamics over time (Fig. 2c, S1). This shows that the structure of the pore remains roughly rotationally symmetric throughout the transition. It is important to note that this barrier-less type of behavior is not guaranteed when using these models; the reason is that, even if both the potential energy and the free energy along the reaction coordinate have a single minimum near the native state (like in the case for some small proteins), kinetic bottlenecks can still be observed, as the effective diffusion coefficient of the protein folding process is usually a function of the reaction coordinate and barriers further impact the lifetime of the intermediates.^40^ In addition, using similar models, the folding of small proteins and conformational changes in molecular complexes are often associated with large-scale rate-limiting free-energy barriers.^63–65^ When monitoring the diameter, it appears that neither effect is significant for constriction of the NPC. Nevertheless, *local* kinetic barriers are present in the system. For example, the linker NUP35 sometimes blocks a small region of the latter, delaying completion of the conformational transition of NUP155 (Fig. S2). However, these are signatures of local kinetic barriers, and they do not manifest as barriers along the global reaction coordinates. Intermediates with long lifetimes in large complexes can often be related to the biological function.^66,67^ Since these types of intermediates appear to be absent in NPC, it is possible that the complex has evolved to be very responsive to external forces introduced by the membrane. That is, the lack of clear intermediates suggests that changes in cellular conditions may lead to the adoption of a continuous spectrum of dilation states, rather than there being an all-or-nothing response.

The second major feature is that the constriction timescales of the outer rings (CR and NR) differ significantly from those of the inner layer of IR and LR (Fig. 2b, c, S1). Since the specific timescales are highly reproducible between simulations, the separation of timescales is statistically significant. This distinction in the constriction times suggests that the CR and NR may undergo rapid reorganization, possibly because they have larger interfaces between spokes in the initial dilated state compared with IR (see the dilated states in Fig. 1d, e). Although the LR is anchored on the IR only by the transmembrane and linker NUPs, it appears the constriction of IR and LR is cooperative. We also infer that movements in the outer rings (CR and NR) are weakly coupled to those of IR and LR. To test this notion, we constructed a perturbed Hamiltonian and repeated our simulations. Structurally, all the forces from the CR and NR are exerted on IR via the Bridge NUP155, which is the only structural element connecting them. By altering the Hamiltonian, such that all the terms related to the Bridges have the same energy in the dilated and constricted states, we define the Bridges to be structurally agnostic and unbiased towards either state. In this sense, the constriction of the CR/NR and IR/LR is decoupled with this Hamiltonian. With the “decoupled” Hamiltonian, the constriction of IR is minimally affected (10%), suggesting that there is a low degree of coupling in the original Hamiltonian (Fig. S3). That is, while the elements are structurally connected, the structure of the NPC rings are intrinsically “weak” in terms of mechanical coupling.

### C. Progressive ordering of the inter-spoke interfaces

Although the system undergoes a constriction of ~ 100 Å, the interactions and forces are short-range. As described above, the structures suggest that IR constriction is likely controlled by relatively local changes in the linkers (NUP35 in Fig. 1d).^8^ This inspired us to study the physical characteristics of these molecules as they transit between states. In this section, we discuss the apparent behavior of each region during constriction, mainly from the perspective of local interactions and local reaction coordinates.

Before discussing statistical descriptions, it is illustrative to first consider the dynamics of a representative simulation, with a focus on a single pair of adjacent spokes. For this, we selected all the NUPs with an RMSD between the dilated and constricted states greater than 5 Å, and analyzed the RMSD change throughout the constriction process. For IR, the selection includes 2 copies of NUP155 (Fig.1c) and 4 copies of NUP35 (Fig.1c,d, 3b). NUP98 in IR also contains a disordered subsequence that forms the diffusion barrier in the central channel of the pore, but this region is not included in either this RMSD analysis or the simulations; its effect on the human NPC is implicitly expressed in the coarse-grained Hamiltonian. The heads of the linker NUPs in adjacent IR spokes interact with each other,^7^ and we can group four NUP35 molecules (two head-to-head pairs) at each inter-spoke interface (Fig. 3b). The individual RMSD values of the linker NUPs at the same interface exhibit strong correlations, as the conformations of all the linkers at a given interface reach the steady state at approximately the same time, indicating the formation of new contacts in the inter-spoke IR interface (represented by balls in Fig. 3a, b). This is further evidenced by the reaction coordinate *Q*, the fraction of formed contacts. In this particular kinetic process, we used a denoised *Q* to reduce the effect of thermal fluctuations (see Methods). A sharp transition in *Q*_interface_ suggests that new contacts that are unique to the constricted state are formed at the inter-spoke IR interface within a short time period as the conformations of the linkers reach a steady state (Fig. 3c, d). Interestingly, the eight inter-spoke IR interfaces always form asynchronously (Fig. 3d, e), despite the symmetry in the structure and force field, as well as the roughly symmetric dynamics in the global reaction coordinate *d*_IR_ (Fig. 2c). For completeness, there are some points about the structure of the NPC that are worth mentioning. The last 20 residues of the linker NUP35 are excluded from the RMSD calculation. This region contains the amphipathic alpha helix tail that interacts with the membrane,^7^ and the Hamiltonian does not stabilize the amphipathic alpha helix tail of Linker 4. The nuclear membrane is not explicitly present in the simulation, though the interactions are phenomenologically integrated in the Hamiltonian as Lennard-Jones potentials for native contacts. However, the fluctuation of the amphipathic helix is more likely an artifact from the construction of the constricted state rather than due to the absence of the membrane, because the same helices from the other three linkers, in a similar environment, are stabilized. The other NUP in the IR with an RMSD larger than 5 Å is NUP155, which normally drops below 3 Å within 250 *t*_0_, indicating a smaller effect on the IR constriction compared with the linker NUPs. One exception is that NUP35 sometimes blocks a small region of NUP155 and delays its full transition (Fig. S2), but this should have little effect on the overall transition.

**FIG. 3.**
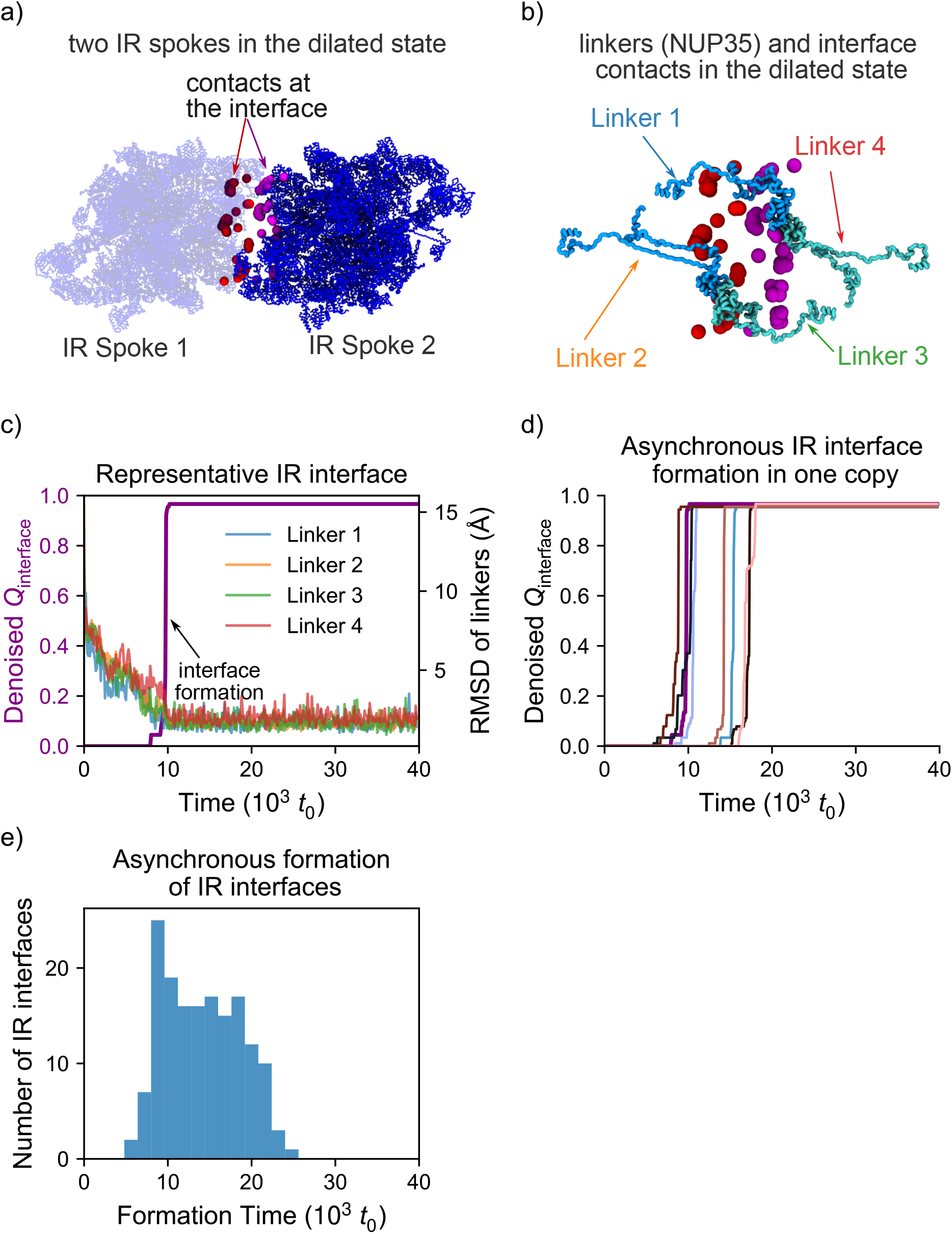
Inter-spoke IR interface formation is asynchronous and abrupt. a) The inter-spoke IR interface between two IR spokes, slightly rotated compared with Fig. 1d. The C_*α*_ atoms involved in interface contacts are colored red and purple for the two different spokes. b) Zoomed-in view of the interface between two adjacent IR spokes. The linkers (NUP35) are colored in light blue and cyan for the two different spokes and are labeled as in panel c). c) Interface formation between IR spokes for a representative trajectory. The individual linker RMSDs reach a steady state at the same time that the inter-spoke contacts are formed. *Q* is measured for the contacts highlighted in panels a) and b). d) The formation of all eight IR interfaces in a representative simulation copy is asynchronous. The thick purple line represents the interface in panel c). e) The distribution of formation times among all interfaces and simulations exhibits asynchronous interface formation.

The outer rings CR and NR are structurally similar, with many components in common. The NUPs with the largest RMSD (one NUP93 and two NUP133s) changes are located at the inter-spoke interfaces (Fig. 1c, e, S5). These NUPs consist of ordered motifs in which the inter-motif distance is adjusted by the disordered motif in between. Taking CR as an example, we also observed a gradual RMSD decrease from the aforementioned NUPs that is correlated with the inter-spoke interface formation, which also remains asynchronous among interfaces (Fig. S5). However, the changes in the NUP conformation, as well as the reaction coordinates *Q*_interface_ and *d*_CR_ are approximately one order of magnitude faster in the simulations than in IR and LR. Another distinction from the IR is that new contacts are formed between spokes at the beginning of the simulations for CR, possibly because the adjacent spokes are already in contact in the initial dilated state (Fig. 1e, S5). For NR, its inter-spoke interfaces are less protected than those of CR, and one of the NUP133 copies is likely to detach at some interfaces, without affecting the NR constriction time. This suggests again that the local asynchrony in interface formation does not affect the symmetry in the global reaction coordinate *d*.

Most NUPs in LR and all the Bridge NUPs have RMSD greater than 5 Å between the dilated and constricted states. The LR of human NPC includes eight copies of NUP210, residing between the two nuclear membrane layers, separated from the IR by the fused region of the membrane. The trans-membrane proteins NDC1 and Aladin, which remain largely unchanged during the transition, are also classified in LR. An asymmetrical unit is formed by eight NUP210s that connect to their counterparts in the adjacent spokes, forming a ring structure among the spokes; the entire ring is connected to the IR via the transmembrane proteins (Fig. 1f). For LR, there is no obvious interface formation during constriction, and all the NUP210s undergo a large conformational change during the constriction with large fluctuations, where the timescale of the RMSD change is comparable to the overall diameter decrease (Fig. S4). More specifically, the bending motion of all the NUP210s narrows the spreading angle of the LR spoke, enabling a decrease in diameter (Fig.1f). For Bridges, they gradually switch conformations to accommodate the asynchronous constriction between the CR/NR and IR (Fig. S4).

### D. Intra-linker interactions drive IR constriction

To complete our analysis of constriction, we asked whether there are specific sets of interactions that more strongly influence the dynamics, especially the slow constriction of IR. For this, we compared the kinetics when different sets of IR contacts were scaled by 0.1. At a qualitative level, this form of analysis is analogous to *φ* -value analysis for protein folding. In *φ* -value analysis, point mutations are introduced to determine whether specific interactions are formed before, or after, the system has crossed the rate-limiting free-energy barrier.^34,68–70^ Here, we adopt the strategy of perturbing sets of IR contacts, to obtain similar insights. For the NPC, while we do not have evidence of a large-scale free-energy barrier, comparison of kinetics can indicate the degree to which different types of interactions shift the energy landscape. For this discussion, the original Hamiltonian described above will be referred to as *H*_0_, and perturbed Hamiltonians are labeled *H*_1_-*H*_4_. In the perturbed Hamiltonians, contacts that are specific to the constricted conformation were weakened. In *H*_1_, intra-linker contacts were weakened (Fig. 4d). In *H*_2_, intramolecular non-linker IR contacts are weaker (Fig. 4e). In *H*_3_, inter-molecular linker contacts were perturbed (Fig. 4f) and in *H*_4_ the inter-spoke IR interfaces were scaled (Fig. 4g).

**FIG. 4.**
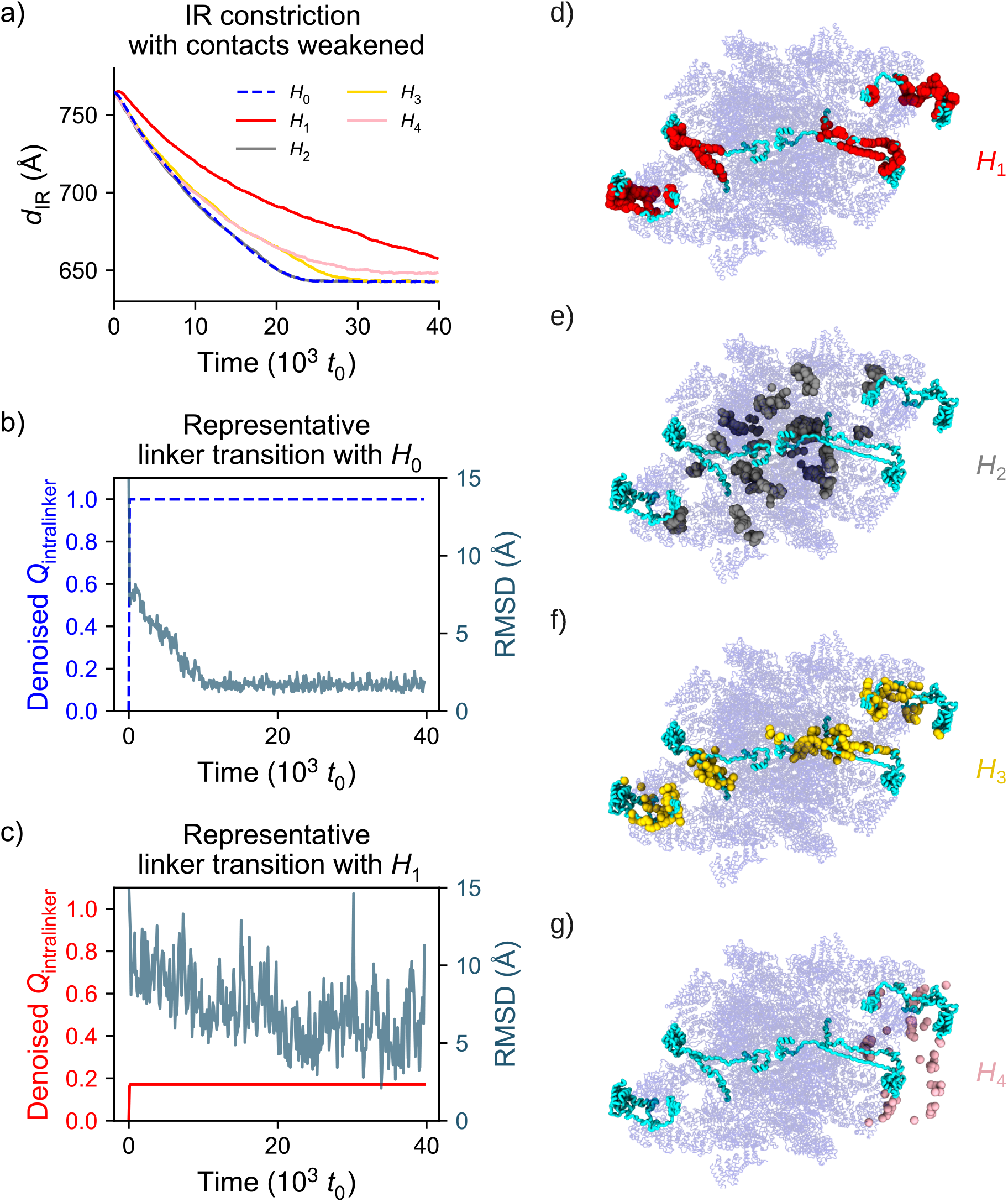
IR constriction is strongly influenced by contact formation in the linkers (NUP35). a) The average IR diameter for variants of the model. Blue dashed line denotes *H*_0_. In *H*_1_ (red), the intra-linkers contacts are scaled by 0.1. For *H*_2_ (grey), the intramolecular non-linker IR contacts are scaled by 0.1. For *H*_3_ (gold), the intermolecular linker contacts are scaled by 0.1. In *H*_4_ (pink), the inter-spoke IR interface contacts are scaled by 0.1. b) The conformational change of a representative linker for *H*_0_. The intra-linker contacts are formed in the early stage of the transition, while the conformational change, as measured by RMSD of NUP35, occurs on a much longer time scale. This suggests that linker contacts formation can lead to geometric frustration that is gradually relaxed during the transition. c) Representative trajectory when using *H*_1_. Most of the intra-linker contacts do no form, the linker conformation remains distant from the constricted structure, and constriction is slower. RMSD of Linker 4 (See Fig. 3b) is shown in both panels b) and c). d-g) The atoms involved in the weakened contacts for *H*_1_, *H*_2_, *H*_3_, and *H*_4_. IR is shown in the dilated state and the four copies of NUP35 are shown in cyan. Contacts in one spoke are shown in panels d) to f), and the contacts in one inter-spoke interface are shown in panel g).

We first characterize *H*_1_ and *H*_2_ to illustrate the effects of intramolecular contacts on IR constriction (in Fig. 4a, d, e, S6). Based on analysis in Section III C, contacts within the linkers are expected to contribute more than the other NUPs to IR constriction. Consistent with this, we find that the constriction timescale difference is most significantly slowed in *H*_1_ (intra-linker perturbation). The timescale observed with *H*_1_ is double that of the original model, while the overall timescale is almost identical for *H*_2_ (Fig. 4a, S6). Interestingly, the number of contacts adjusted in *H*_1_ and *H*_2_ only differs by 28%. Accordingly, one may not explain the effect solely based on the number of contacts within each group. Next, we investigated the effects of the interactions between linkers and the other NUPs in the IR with *H*_3_ (Fig. 4a, f, S6). The results show that, when the intermolecular contacts involving the linkers are weakened, there is a more modest (∼ 30%) reduction in kinetics, further reinforcing the central role of intra-linker interactions. We then probed the effects of the IR-interfaces using *H*_4_ (Fig. 4a, g, S6). This Hamiltonian yields results similar to *H*_3_, except that the final state features a larger diameter. This indicates that the interfaces are not critical to constriction kinetics, though they do facilitate final rearrangements within nearly-fully-constricted states.

Comparison of the timing of contact formation and the effect on kinetics also provides a description of how the NPC may accumulate and release strain energy during constriction/dilation. For example, the vast majority of intra-linker contacts form within 100 *t*_0_ with *H*_0_ (Fig. 4b). However, formation of contacts is not associated with immediate contraction of the linkers. Rather, the linkers undergo gradual changes and both the linker NUP conformational change and the IR constriction occur on much longer timescales (two orders of magnitude longer). This may imply that early contact formation introduces local geometric frustration that ultimately drives the constriction process. That is, the structure of the NPC cannot simultaneously satisfy all the energetically favored local interactions,^71^ resulting in some local conformations in the high-energy state that can only be resolved through conformational changes on larger scales. This interpretation is also consistent with the apparent large structural deviations of the linkers when using Hamiltonian *H*_1_ (Fig. 4c). In terms of future efforts, these observations suggest that more precise analysis of linker energetics will be central to establishing a comprehensive understanding of how specific interactions contribute to the overall constriction process.

One interesting prediction is that different sets of interactions contribute to distinct stages of the constriction process. That is, when intra-linker interactions are changed (*H*_1_), the NPC constriction is delayed, even at short times. However, in contrast, for *H*_3_ and *H*_4_, the delay in constriction only becomes evident at longer timescales, at which point approximately half of the constriction process has completed. Further, there is an additional delay in constriction for *H*_4_ (interface perturbation), consistent with the interfaces forming late in the constriction process.

## V. DISCUSSION

The human NPC responds to external forces from the nuclear membrane. This adaptation itself is passive but is coupled to active processes that change the nuclear size. We consider two extreme states observed in experiments and the transition pathway between them. All the external forces are incorporated in the SBM Hamiltonian, which is a function of only the C_*α*_ atom positions of the human NPC. Equivalently, the human NPC is initially equilibrated at the dilated state with effective external forces from the membrane. With the external forces altered at *t* = 0, the NPC undergoes relaxation toward a new equilibrium near the constricted state. This approach allows us to probe both the global feature of NPC dynamics and the contribution from local structural units.

Different rings of the human NPC exhibit distinct timescales from the constriction simulations. The structurally similar CR and NR both undergo a relatively fast constriction, while the IR, whose diameter change is only weakly coupled to those of CR and NR, follows a similar pattern but on a much longer timescale. The global conformational change of each ring can be connected to a few NUPs at the inter-spoke interfaces. For CR and NR, three NUPs are identified at the inter-spoke interface, with large conformational changes associated with local disordered regions, and a transition time comparable to that of CR constriction. IR is of the most interest among all the regions of the NPC, as it anchors the transport barrier proteins in the central channel. Given the rigidity of the scaffold structure of the IR spokes, the IR constriction is largely affected by the conformational change in the linker NUP35. This type of NUP consists of ordered regions that Bridge the two IR spokes and disordered regions that are tethered to the remainder of the IR spoke. They undergo a slow and asynchronous conformational change, drive the IR spokes and close the gap between them, and eventually pull the IR spokes close to each other.

The motion of IR is coupled to the other rings by various means in our force field. The LR is connected to the IR via the cross-membrane protein, whose conformational change during constriction is negligible. The diameter of LR is strongly correlated with the IR, where NUP210 of the LR bends to accommodate the reduced diameter. The CR and NR are connected to the IR via the Bridge NUP155. As the CR and NR relax to the constricted state earlier than the IR, the Bridges have to accommodate the constricted state of the CR and NR, and the (almost) dilated state of the IR. However, this potentially frustrated conformation has only a small contribution to IR constriction, indicating that the Bridges are sufficiently flexible to accommodate different ring states.

The interplay between the nuclear membrane and the NPC is implicitly expressed in the Hamiltonian, of which a more comprehensive understanding may benefit from further work. It is suggested that NUP160 and NUP133 of the CR and NR, the Bridge NUP155, the linker NUP35, the NUP155 copy belonging to IR, along with the transmembrane proteins, are anchored on the nuclear membrane.^7^ The NUP155, NUP35, and the transmembrane proteins form a transmembrane interaction hub. The motion of CR, NR, and IR can then be coupled through their interaction with the membrane. One may also speculate that the membrane exerts force on the linkers and NUP133 in the outer rings *in cellulo*, driving the conformational change that prevents IR interface formation and modulates the transport barrier in the center. While many aspect of NPC dynamics have yet to be probed, the current study provides a framework for precise characterization of the energetic and structural factors that dynamically regulate the NPC.

## V. ACKNOWLEDGEMENTS

We thank Peter Wolynes for useful discussions on protein dynamics and frustration. This research was supported by the NSF (PHY-2014141, and PHY-2210291), NIH (Grant R35GM153502-02) and Welch Foundation (Grant C-1792). Work at the Center for Theoretical Biological Physics was supported by the NSF (Grant PHY-2019745). VGC also acknowledges support from the NSF Award PHY-2609969 and the NVIDIA Academic Grant Program. JNO is a Cancer Prevention and Research Institute of Texas (CPRIT) Scholar in Cancer Research. Financial support was also provided in part by the Brazilian agencies Fundação de Amparo à Pesquisa do Estado de Minas Gerais (FAPEMIG, APQ-03197-24) and Conselho Nacional de Desenvolvimento Científico e Tecnológico (CNPq, 307992/2022-5). We would like to thank AMD (Advanced Micro Devices, Inc.) for donating critical hardware and support resources from its HPC Fund, which made this work possible.

## VI. DATA AVAILABILITY

The data that support the findings of this study are available on reasonable request from the corresponding author.

