## Supporting Information for "Asynchronous structural rearrangements govern constriction of the human nuclear pore complex"

### I. LIST OF FIGURES

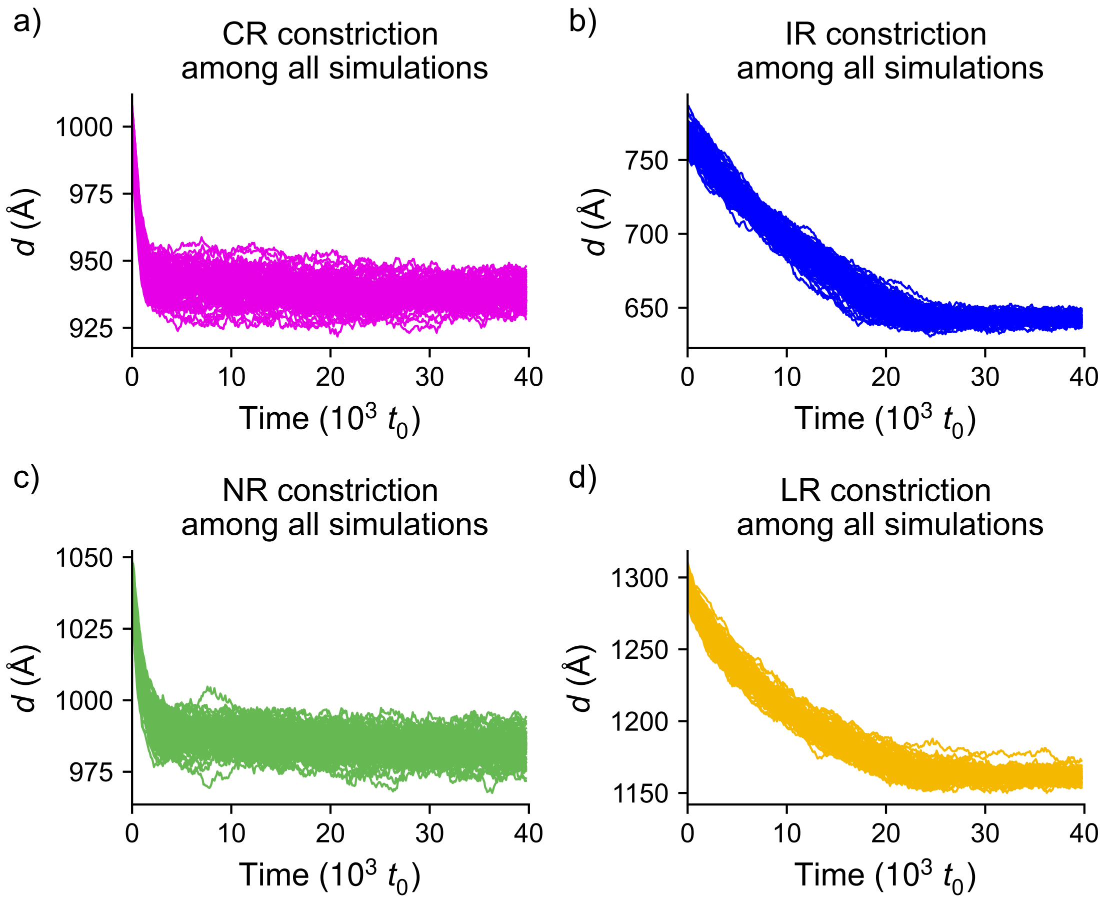


FIG. S1. Diameter change of a) CR, b) IR, c) NR, d) LR for all 20 simulated events (shown in Fig. 2c). As the NPC has 8 spokes, 4 diameters are measured, yielding 80 trajectories in the diameter space.

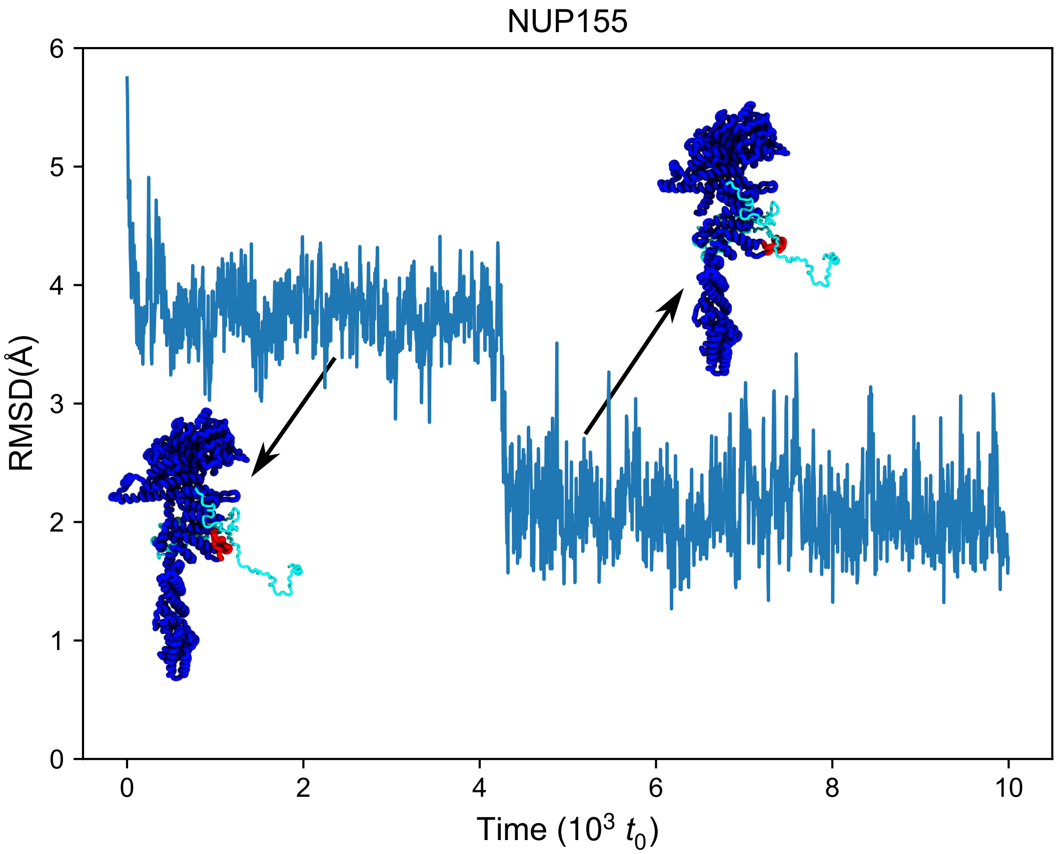


FIG. S2. One of NUP155 proteins in IR (four in an IR spoke) is likely to get trapped by the linker NUP (cyan). The RMSD of the trapped NUP155 compared to the constricted state exhibits a $4 \times 10^3$ $t_0$ plateau where a few residues (red) from NUP155 are sterically blocked by the linker NUP.

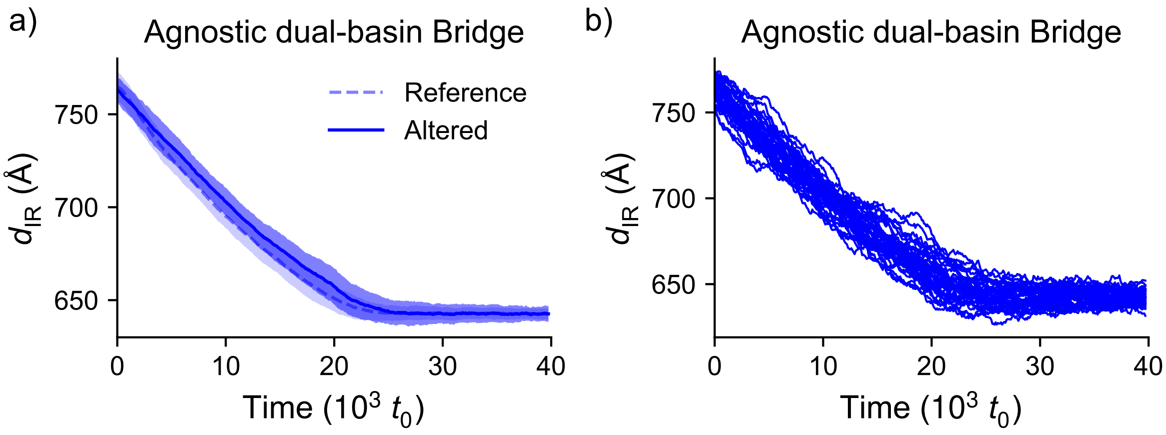


FIG. S3. The Hamiltonian with agnostic dual-basin Bridges shows a similar IR constriction pattern as the original Hamiltonian. Specifically, the perturbed Hamiltonian has a balanced energy in the dilated and constricted states for all Bridges between CR/NR and IR. a) The average and sample standard deviation of the IR diameters during the transition. The diameters evolve similarly between the original and perturbed Hamiltonians. b) The original IR diameter data corresponding to the perturbed Hamiltonian in panel a), acquired from 10 simulations, for a total of 40 trajectories in the diameter space.

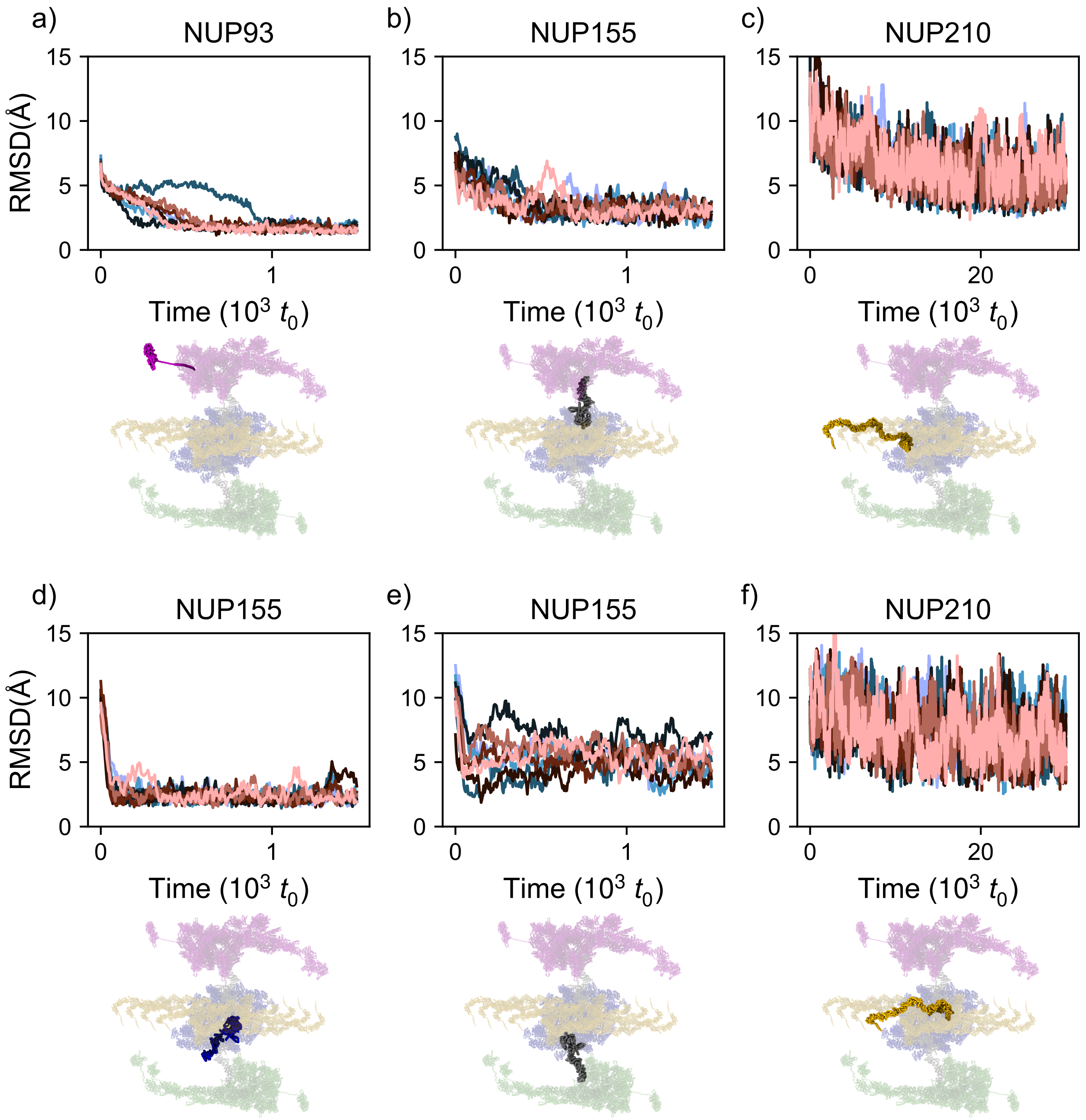


FIG. S4. For the NUPs exhibiting substantial conformational differences between the dilated and constricted states (excluding NUP210), their RMSD relative to the constricted state decreases progressively in the simulation, with the conformations eventually fluctuating around the constricted structure. Panels a) to f) present six NUPs as examples. For each NUP, the upper panel shows the RMSD of the corresponding protein in all eight spokes during a representative simulation, and the lower panel shows its location within the NPC.

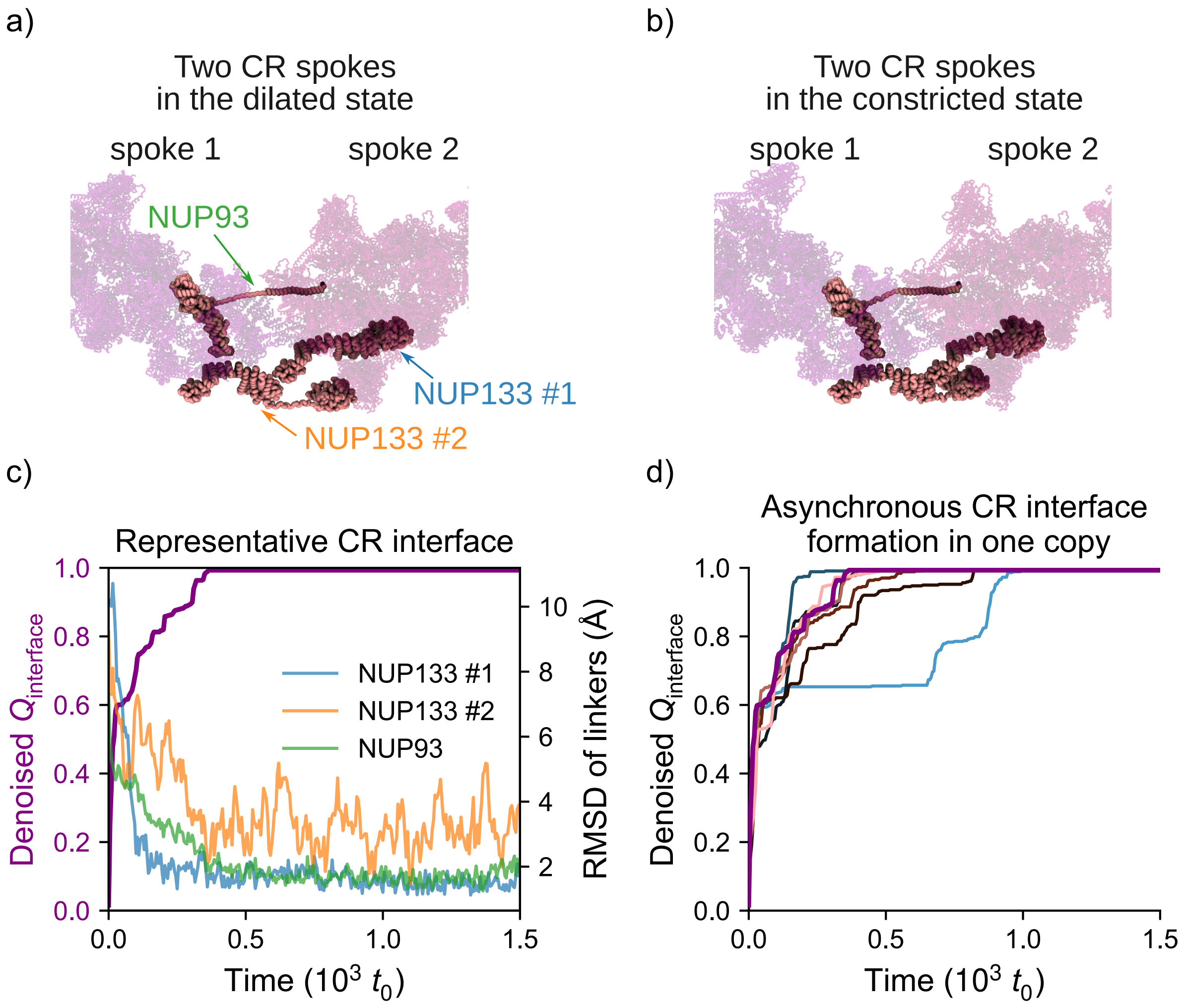


FIG. S5. Inter-spoke CR interface formation is asynchronous. a-b) The interface between two adjacent CR spokes in a) dilated and b) constricted states, zoomed in from Fig. 1e. The NUPs whose RMSD between the two states is greater than 5 Å (one NUP93 and two NUP133s) are colored in pink, and all the other NUPs are shown transparently. c) A representative interface formation event between CR spokes. The NUPs are colored the same as their labels in panel a). d) The formation of the eight CR interfaces in a representative simulation. The thick purple line represents the interface shown in panel c).

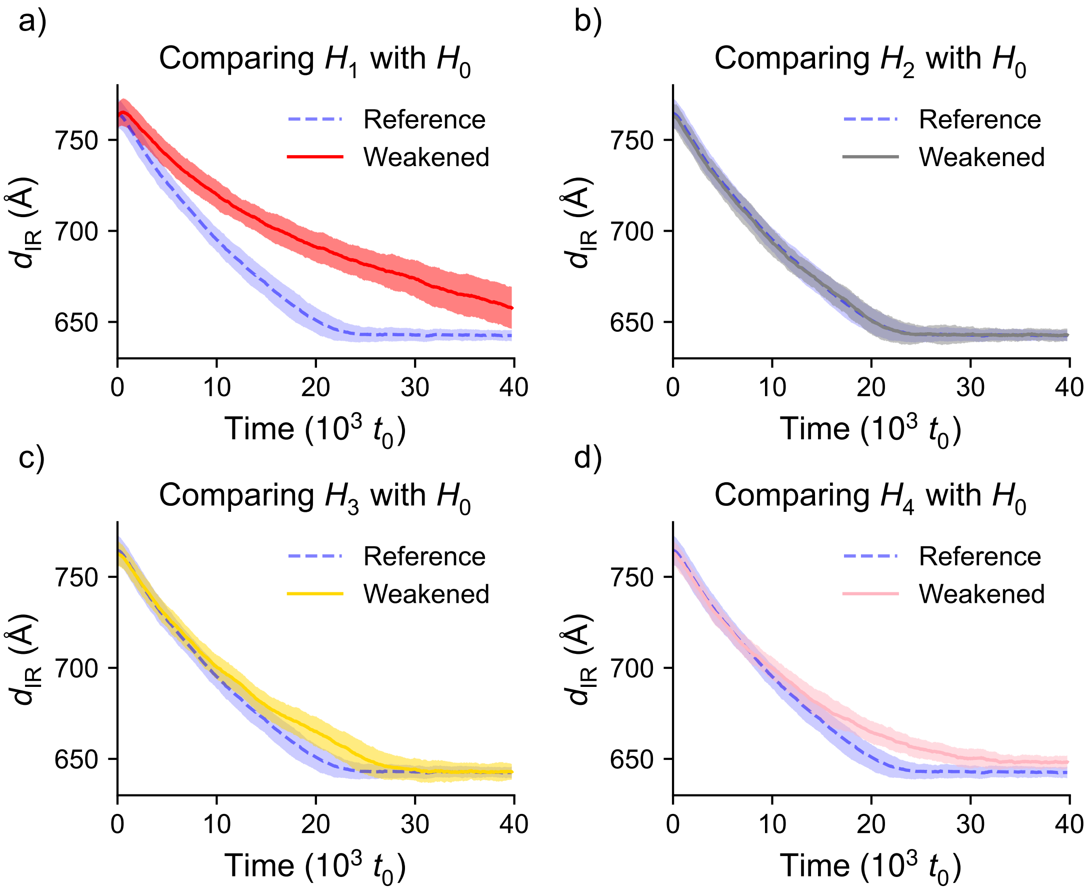


FIG. S6. IR diameter as a function of time for the original Hamiltonian ($H_0$) and a) $H_1$, b) $H_2$, c) $H_3$, d) $H_4$. Dashed and solid lines represent the average IR diameter (the same data shown in Fig. 4a) with the original and perturbed Hamiltonians, respectively, and shaded regions denote the sample standard deviation.
